## Supplementary material for "Workflow sharing with automated metadata validation and test execution to improve the reusability of published workflows": LaTeX Source: api-response.pdf

```
$ curl -fsSL https://pitagora-network.org/yevis-DAT2-cwl/tools/d03458d8-837c-4173-afa3-
55ebe538b0b2/versions/1.0.0 | jq .
{
  "author": [
    "suecharo"
  ],
  "name": "DAT2-cwl - bacteria genome workflow",
  "url": "https://pitagora-network.github.io/yevis-DAT2-cwl/tools/d03458d8-837c-4173-afa3-
55ebe538b0b2/versions/1.0.0",
  "id": "1.0.0",
  "descriptor_type": [
    "CWL"
  ],
  "verified": true,
  "verified_source": [
    "https://github.com/pitagora-network/yevis-DAT2-cwl/actions/runs/2317749577"
  ]
}
```
