## Supplementary material for "Workflow sharing with automated metadata validation and test execution to improve the reusability of published workflows": LaTeX Source: main.bib

@article{goodwin_coming_2016,
  title        = {{Coming of age: Ten years of next-generation sequencing technologies}},
  author       = {Goodwin, Sara and {McPherson}, John D. and {McCombie}, Richard W.},
  year         = 2016,
  journal      = {{Nature Reviews Genetics}},
  volume       = 17,
  number       = 6,
  pages        = {333--351},
  doi          = {10.1038/nrg.2016.49}
}
@article{stein_case_2010,
  title        = {{The case for cloud computing in genome informatics}},
  author       = {Stein, Lincoln D.},
  year         = 2010,
  journal      = {{Genome Biology}},
  volume       = 11,
  number       = 5,
  pages        = 207,
  doi          = {10.1186/gb-2010-11-5-207}
}
@article{prins_toward_2015,
  title        = {{Toward effective software solutions for big biology}},
  author       = {Prins, Pjotr and de Ligt, Joep and Tarasov, Artem and Jansen, Ritsert C. and Cuppen, Edwin and Bourne, Philip E.},
  year         = 2015,
  journal      = {{Nature Biotechnology}},
  volume       = 33,
  number       = 7,
  pages        = {686--687},
  doi          = {10.1038/nbt.3240}
}
@article{perkel_workflow_2019,
  title        = {{Workflow systems turn raw data into scientific knowledge}},
  author       = {Perkel, Jeffrey M.},
  year         = 2019,
  journal      = {{Nature}},
  volume       = 573,
  number       = 7772,
  pages        = {149--150},
  doi          = {10.1038/d41586-019-02619-z}
}
@article{leprevost_best_2014,
  title        = {{On best practices in the development of bioinformatics software}},
  author       = {Leprevost, Felipe da Veiga and Barbosa, Valmir C. and Francisco, Eduardo L. and Perez-Riverol, Yasset and Carvalho, Paulo C.},
  year         = 2014,
  journal      = {{Frontiers in Genetics}},
  volume       = 5,
  doi          = {10.3389/fgene.2014.00199}
}
@article{afgan_galaxy_2018,
  title        = {{The Galaxy platform for accessible, reproducible and collaborative biomedical analyses: 2018 update}},
  author       = {Afgan, Enis and Baker, Dannon and Batut, B\'{e}r\'{e}nice and van~den~Beek, Marius and Bouvier, Dave and \v{C}ech, Martin and Chilton, John and others},
  year         = 2018,
  journal      = {{Nucleic Acids Research}},
  volume       = 46,
  pages        = {W537--W544},
  doi          = {10.1093/nar/gky379},
  issue        = {W1}
}
@article{crusoe_methods_2021,
  title        = {{Methods included: Standardizing computational reuse and portability with the common workflow language}},
  author       = {Crusoe, Michael R. and Abeln, Sanne and Iosup, Alexandru and Amstutz, Peter and Chilton, John and Tijani\'{c}, Neboj\v{s}a and M\'{e}nager, Herv\'{e} and others},
  year         = 2021,
  journal      = {{arXiv}},
  doi          = {10.48550/arXiv.2105.07028},
  url          = {https://doi.org/10.48550/arXiv.2105.07028}
}
@article{voss_full-stack_2017,
  title        = {{Full-stack genomics pipelining with GATK4 + WDL + Cromwell}},
  author       = {Voss, Kate and Gentry, Jeff and Auwera, Geraldine Van Der},
  year         = 2017,
  journal      = {{F1000Research}},
  doi          = {10.7490/F1000RESEARCH.1114631.1}
}
@article{di_tommaso_nextflow_2017,
  title        = {{Nextflow enables reproducible computational workflows}},
  author       = {Di Tommaso, Paolo and Chatzou, Maria and Floden, Evan W. and Barja, Pablo Prieto and Palumbo, Emilio and Notredame, Cedric},
  year         = 2017,
  journal      = {{Nature Biotechnology}},
  volume       = 35,
  number       = 4,
  pages        = {316--319},
  doi          = {10.1038/nbt.3820}
}
@article{koster_snakemake_2012,
  title        = {{Snakemake---a scalable bioinformatics workflow engine}},
  author       = {Koster, J. and Rahmann, S.},
  year         = 2012,
  journal      = {{Bioinformatics}},
  volume       = 28,
  number       = 19,
  pages        = {2520--2522},
  doi          = {10.1093/bioinformatics/bts480}
}
@article{da_veiga_leprevost_biocontainers_2017,
  title        = {{BioContainers: An open-source and community-driven framework for software standardization}},
  author       = {Leprevost, Felipe da Veiga and Gr\"{u}ning, Bj\"{o}rn A. and Alves Aflitos, Saulo and R\"{o}st, Hannes L. and Uszkoreit, Julian and Barsnes, Harald and Vaudel, Marc and others},
  year         = 2017,
  journal      = {{Bioinformatics}},
  volume       = 33,
  number       = 16,
  pages        = {2580--2582},
  doi          = {10.1093/bioinformatics/btx192},
  editor       = {Valencia, Alfonso}
}
@article{goble_implementing_2021,
  title        = {{Implementing FAIR Digital Objects in the EOSC-Life Workflow Collaboratory}},
  author       = {Goble, Carole and Soiland-Reyes, Stian and Bacall, Finn and Owen, Stuart and Williams, Alan and Eguinoa, Ignacio and Droesbeke, Bert and others},
  year         = 2021,
  journal      = {{Zenodo}},
  publisher    = {Zenodo},
  doi          = {10.5281/zenodo.4605654}
}
@article{oconnor_dockstore_2017,
  title        = {{The Dockstore: Enabling modular, community-focused sharing of Docker-based genomics tools and workflows}},
  author       = {O'Connor, Brian D. and Yuen, Denis and Chung, Vincent and Duncan, Andrew G. and Liu, Xiang Kun and Patricia, Janice and Paten, Benedict and others},
  year         = 2017,
  journal      = {{F1000Research}},
  volume       = 6,
  pages        = 52,
  doi          = {10.12688/f1000research.10137.1}
}
@article{ewels_nf-core_2020,
  title        = {{The nf-core framework for community-curated bioinformatics pipelines}},
  author       = {Ewels, Philip A. and Peltzer, Alexander and Fillinger, Sven and Patel, Harshil and Alneberg, Johannes and Wilm, Andreas and Garcia, Maxime Ulysse and others},
  year         = 2020,
  journal      = {{Nature Biotechnology}},
  volume       = 38,
  number       = 3,
  pages        = {276--278},
  doi          = {10.1038/s41587-020-0439-x}
}
@article{rehm_ga4gh_2021,
  title        = {{GA4GH: International policies and standards for data sharing across genomic research and healthcare}},
  author       = {Rehm, Heidi L. and Page, Angela J. H. and Smith, Lindsay and Adams, Jeremy B. and Alterovitz, Gil and Babb, Lawrence J. and Barkley, Maxmillian P. and others},
  year         = 2021,
  journal      = {{Cell Genomics}},
  volume       = 1,
  number       = 2,
  pages        = 100029,
  doi          = {10.1016/j.xgen.2021.100029}
}
@article{wilkinson_fair_2016,
  title        = {{The FAIR Guiding Principles for scientific data management and stewardship}},
  author       = {Wilkinson, Mark D. and Dumontier, Michel and Aalbersberg, {IJsbrand} Jan and Appleton, Gabrielle and Axton, Myles and Baak, Arie and Blomberg, Niklas and others},
  year         = 2016,
  journal      = {{Scientific Data}},
  volume       = 3,
  number       = 1,
  pages        = 160018,
  doi          = {10.1038/sdata.2016.18}
}
@article{khan_sharing_2019,
  title        = {{Sharing interoperable workflow provenance: A review of best practices and their practical application in CWLProv}},
  author       = {Khan, Farah Zaib and Soiland-Reyes, Stian and Sinnott, Richard O. and Lonie, Andrew and Goble, Carole and Crusoe, Michael R.},
  year         = 2019,
  journal      = {{GigaScience}},
  volume       = 8,
  number       = 11,
  pages        = {giz095},
  doi          = {10.1093/gigascience/giz095}
}
@article{broad_sequencing_platform_members_genome_2005,
  title        = {{Genome sequence, comparative analysis and haplotype structure of the domestic dog}},
  author       = {Lindblad-Toh, Kerstin and Wade, Claire M and Mikkelsen, Tarjei S. and Karlsson, Elinor K. and Jaffe, David B. and Kamal, Michael and Clamp, Michele and others},
  year         = 2005,
  journal      = {{Nature}},
  volume       = 438,
  number       = 7069,
  pages        = {803--819},
  doi          = {10.1038/nature04338}
}
@article{loman_complete_2015,
  title        = {{A complete bacterial genome assembled de novo using only nanopore sequencing data}},
  author       = {Loman, Nicholas J and Quick, Joshua and Simpson, Jared T},
  year         = 2015,
  journal      = {{Nature Methods}},
  volume       = 12,
  number       = 8,
  pages        = {733--735},
  doi          = {10.1038/nmeth.3444}
}
@article{merkel_docker_2014,
  title        = {{Docker: Lightweight Linux containers for consistent development and deployment}},
  author       = {Merkel, Dirk},
  year         = 2014,
  journal      = {{Linux Journal}},
  volume       = 2014,
  pages        = 2
}
@book{bono_dat2_2019,
  title        = {{Next Generation Sequencer DRY Analysis Manual 2nd Edition}},
  author       = {Hidemasa, Bono and Shimizu, Atsushi},
  year         = 2019,
  publisher    = {{Gakken Medical Shujunsha}}
}
@article{shen_seqkit_2016,
  title        = {{SeqKit: A cross-platform and ultrafast toolkit for FASTA/Q file manipulation}},
  author       = {Shen, Wei and Le, Shuai and Li, Yan and Hu, Fuquan},
  year         = 2016,
  journal      = {{PLOS One}},
  volume       = 11,
  number       = 10,
  pages        = {e0163962},
  doi          = {10.1371/journal.pone.0163962},
  editor       = {Zou, Quan}
}
@misc{simon_fastqc_2010,
  title        = {{FastQC: A quality control tool for high throughput sequence data}},
  author       = {Andrews, Simon},
  year         = 2010,
  publisher    = {{Babraham Institute}},
  url          = {https://www.bioinformatics.babraham.ac.uk/projects/fastqc/}
}
@article{chen_fastp_2018,
  title        = {{fastp: An ultra-fast all-in-one FASTQ preprocessor}},
  author       = {Chen, Shifu and Zhou, Yanqing and Chen, Yaru and Gu, Jia},
  year         = 2018,
  journal      = {{Bioinformatics}},
  volume       = 34,
  number       = 17,
  pages        = {i884--i890},
  doi          = {10.1093/bioinformatics/bty560}
}
@article{kajitani_efficient_2014,
  title        = {{Efficient de novo assembly of highly heterozygous genomes from whole-genome shotgun short reads}},
  author       = {Kajitani, Rei and Toshimoto, Kouta and Noguchi, Hideki and Toyoda, Atsushi and Ogura, Yoshitoshi and Okuno, Miki and Yabana, Mitsuru and others},
  year         = 2014,
  journal      = {{Genome Research}},
  volume       = 24,
  number       = 8,
  pages        = {1384--1395},
  doi          = {10.1101/gr.170720.113}
}
@misc{ga4gh_wes-api_2017,
  title        = {{ga4gh/workflow-execution-service-schemas}},
  author       = {{Global Alliance for Genomics and Health}},
  year         = 2017,
  publisher    = {GitHub},
  url          = {https://github.com/ga4gh/workflow-execution-service-schemas},
  howpublished = {\url{https://github.com/ga4gh/workflow-execution-service-schemas}}
}
@misc{ga4gh_trs-api_2016,
  title        = {{ga4gh/tool-registry-service-schemas}},
  author       = {{Global Alliance for Genomics and Health}},
  year         = 2016,
  publisher    = {GitHub},
  url          = {https://github.com/ga4gh/tool-registry-service-schemas},
  howpublished = {\url{https://github.com/ga4gh/tool-registry-service-schemas}},
  urldate      = {accessed 2021-12-1}
}
@misc{suetake_yevis-getting-started_2022,
  title        = {{Yevis: Getting Started}},
  author       = {Suetake, Hirotaka and Ohta, Tazro},
  year         = 2022,
  publisher    = {GitHub},
  doi          = {10.5281/zenodo.6793218},
  url          = {https://sapporo-wes.github.io/yevis-cli/getting\_started},
  howpublished = {\url{https://sapporo-wes.github.io/yevis-cli/getting_started}},
  version      = {0.5.1}
}
@article{suetake_sapporo_2022,
	title        = {{Sapporo: A workflow execution service that encourages the reuse of workflows in various languages in bioinformatics}},
	author       = {Suetake, Hirotaka and Tanjo, Tomoya and Ishii, Manabu and P. Kinoshita, Bruno and Fujino, Takeshi and Hachiya, Tsuyoshi and others},
	year         = 2022,
	journal      = {{F1000Research}},
	volume       = 11,
	pages        = 889,
	doi          = {10.12688/f1000research.122924.1}
}
@misc{suetake_yevis-test-action_2022,
  title        = {{ddbj/yevis-cli: 0.5.1 - actions\_example/yevis-test-pr.yml}},
  author       = {Suetake, Hirotaka and Ohta, Tazro},
  year         = 2022,
  publisher    = {GitHub},
  doi          = {10.5281/zenodo.6793218},
  url          = {https://github.com/ddbj/yevis-cli/blob/0.5.1/actions\_example/yevis-test-pr.yml},
  howpublished = {\url{https://github.com/ddbj/yevis-cli/blob/0.5.1/actions_example/yevis-test-pr.yml}},
  version      = {0.5.1}
}
@misc{suetake_yevis-publish-action_2022,
  title        = {{ddbj/yevis-cli: 0.5.1 - actions\_example/yevis-publish-pr.yml}},
  author       = {Suetake, Hirotaka and Ohta, Tazro},
  year         = 2022,
  publisher    = {GitHub},
  doi          = {10.5281/zenodo.6793218},
  url          = {https://github.com/ddbj/yevis-cli/blob/0.5.1/actions\_example/yevis-publish-pr.yml},
  howpublished = {\url{https://github.com/ddbj/yevis-cli/blob/0.5.1/actions_example/yevis-publish-pr.yml}},
  version      = {0.5.1}
}
@misc{pitagora_dat2cwl_2022,
  title        = {{pitagora-network/DAT2-cwl: 1.1.1}},
  author       = {{Pitagora Network members}},
  year         = 2022,
  month        = may,
  publisher    = {Zenodo},
  doi          = {10.5281/zenodo.6565977},
  url          = {https://doi.org/10.5281/zenodo.6565977},
  version      = {1.1.1}
}
@misc{pitagora_bacteria_2022,
  title        = {{GitHub - pitagora-network/DAT2-cwl: 1.1.1 - workflow/bacteria-genome}},
  author       = {{Pitagora Network members}},
  year         = 2022,
  month        = may,
  publisher    = {Zenodo},
  doi          = {10.5281/zenodo.6565977},
  url          = {https://github.com/pitagora-network/DAT2-cwl/tree/1.1.1/workflow/bacteria-genome},
  version      = {1.1.1}
}
@misc{suetake_yevis-dat2-cwl_2022,
  title        = {{pitagora-network/yevis-DAT2-cwl: 1.0.0}},
  author       = {Suetake, Hirotaka},
  year         = 2022,
  publisher    = {Zenodo},
  doi          = {10.5281/zenodo.6572565},
  url          = {https://doi.org/10.5281/zenodo.6572565},
  version      = {1.0.0}
}
@misc{suetake_yevis-metadata-file-bacteria_2022,
  title        = {{Yevis metadata file for the DAT2-cwl/bacteria-genome workflow}},
  author       = {Suetake, Hirotaka},
  year         = 2022,
  month        = may,
  publisher    = {Zenodo},
  doi          = {10.5281/zenodo.6572565},
  url          = {https://github.com/pitagora-network/yevis-DAT2-cwl/blob/1.0.0/d03458d8-837c-4173-afa3-55ebe538b0b2/yevis-metadata-1.0.0.yml},
  version      = {1.0.0}
}
@misc{suetake_yevis-uploaded-files-bacteria_2022,
  title        = {{DAT2-cwl/bacteria-genome workflow files uploaded to Zenodo by Yevis}},
  author       = {Suetake, Hirotaka},
  year         = 2022,
  month        = may,
  publisher    = {Zenodo},
  doi          = {10.5281/zenodo.6545122},
  url          = {https://doi.org/10.5281/zenodo.6545122},
  version      = {1.0.0}
}
@misc{suetake_screenshots_2022,
  title        = {{Screenshots and logs of the DAT2-cwl/bacteria-genome workflow review process and publication process}},
  author       = {Suetake, Hirotaka},
  year         = 2022,
  publisher    = {Zenodo},
  doi          = {10.5281/zenodo.6575319},
  url          = {https://doi.org/10.5281/zenodo.6575319},
  version      = {1.0.0}
}
@misc{suetake_yevis-dat2-cwl-browser_2022,
  title        = {{pitagora-network/yevis-DAT2-cwl-browser: 1.0.0}},
  author       = {Suetake, Hirotaka},
  year         = 2022,
  publisher    = {Zenodo},
  doi          = {10.5281/zenodo.6575089},
  url          = {https://doi.org/10.5281/zenodo.6575089},
  version      = {1.0.0}
}
@misc{suetake_ddbj-workflow-registry_2022,
  title        = {{ddbj/workflow-registry: 1.0.2}},
  author       = {Suetake, Hirotaka and Ohta, Tazro},
  year         = 2022,
  publisher    = {Zenodo},
  doi          = {10.5281/zenodo.6719845},
  url          = {https://doi.org/10.5281/zenodo.6719845},
  version      = {1.0.2}
}
@misc{amstutz2021existing,
  title        = {Existing Workflow systems},
  author       = {Amstutz, Peter and Mikheev, Maxim and Crusoe, Michael R. and Tijani\'{c}, Neboj\v{s}a and Lampa, Samuel},
  year         = 2021,
  howpublished = {\url{https://s.apache.org/existing-workflow-systems}},
  url          = {https://s.apache.org/existing-workflow-systems},
  urldate      = {accessed 2021-12-1}
}
@article{goble2020fair,
  title={{FAIR computational workflows}},
  author={Goble, Carole and Cohen-Boulakia, Sarah and Soiland-Reyes, Stian and Garijo, Daniel and Gil, Yolanda and Crusoe, Michael R. and Peters, Kristian and Schober, Daniel},
  journal={{Data Intelligence}},
  volume={2},
  number={1-2},
  pages={108--121},
  year={2020},
  doi={10.1162/dint_a_00033}
}
@article{wratten_reproducible_2021,
	title        = {{Reproducible, scalable, and shareable analysis pipelines with bioinformatics workflow managers}},
	author       = {Wratten, Laura and Wilm, Andreas and G\"{o}ke, Jonathan},
	year         = 2021,
	journal      = {{Nature Methods}},
	volume       = 18,
	number       = 10,
	pages        = {1161--1168},
	doi          = {10.1038/s41592-021-01254-9}
}
