## Supplementary material for "Workflow sharing with automated metadata validation and test execution to improve the reusability of published workflows": LaTeX Source: main.tex

\documentclass[times, twoside, watermark]{zHenriquesLab-StyleBioRxiv}

\usepackage{blindtext}
\usepackage{graphicx}
\usepackage{siunitx}
\usepackage{flushend}
\usepackage{url}
\usepackage{booktabs}
\usepackage{multirow}
\usepackage{tabularx}

% Please give the surname of the lead author for the running footer
\leadauthor{Suetake}

\begin{document}

\title{Workflow sharing with automated metadata validation and test execution to improve the reusability of published workflows}
\shorttitle{Workflow sharing with automated metadata validation and test execution}

% Use letters for affiliations, numbers to show equal authorship (if applicable) and to indicate the corresponding author
\author[1]{Hirotaka Suetake}
\author[2]{Tsukasa Fukusato}
\author[1]{Takeo Igarashi}
\author[3,\Letter]{Tazro Ohta}

\affil[1]{Department of Creative Informatics, Graduate School of Information Science and Technology, The University of Tokyo, Tokyo, Japan}
\affil[2]{Department of Computer Science, Graduate School of Information Science and Technology, The University of Tokyo, Tokyo, Japan}
\affil[3]{Database Center for Life Science, Joint Support-Center for Data Science Research, Research Organization of Information and Systems, Shizuoka, Japan}

\maketitle

\begin{abstract}
\textbf{\textit{Background:}}
Many open-source workflow systems have made bioinformatics data analysis procedures portable.
Sharing these workflows provides researchers easy access to high-quality analysis methods without the requirement of computational expertise.
However, published workflows are not always guaranteed to be reliably reusable.
Therefore, a system is needed to lower the cost of sharing workflows in a reusable form.

\vspace{1.5mm}
\textbf{\textit{Results:}}
We introduce Yevis, a system to build a workflow registry that automatically validates and tests workflows to be published.
The validation and test are based on the requirements we defined for a workflow being reusable with confidence.
Yevis runs on GitHub and Zenodo and allows workflow hosting without the need of dedicated computing resources.
A Yevis registry accepts workflow registration via a GitHub pull request, followed by an automatic validation and test process for the submitted workflow.
As a proof of concept, we built a registry using Yevis to host workflows from a community to demonstrate how a workflow can be shared while fulfilling the defined requirements.

\vspace{1.5mm}
\textbf{\textit{Conclusions:}}
Yevis helps in the building of a workflow registry to share reusable workflows without requiring extensive human resources.
By following Yevis's workflow-sharing procedure, one can operate a registry while satisfying the reusable workflow criteria.
This system is particularly useful to individuals or communities that want to share workflows but lacks the specific technical expertise to build and maintain a workflow registry from scratch.
\end {abstract}

\begin{keywords}
    Workflow | Workflow language | Continuous integration | Open science | Reproducibility | Reusability
\end{keywords}

\begin{corrauthor}
   
\end{corrauthor}

\section*{Background}
\label{sec:background}

Due to the low cost and high throughput of measurement instruments that acquire digital data from biological samples, the volume of readily available data has become enormous~\citep{goodwin_coming_2016}.
To obtain scientific knowledge from large datasets, a number of computational data analysis processes are required, for example, in DNA sequencing, sequence read trimming, alignment with reference genomes, and annotation using public databases~\citep{stein_case_2010}.
Researchers have developed analysis tools for each process and often publish them as open-source software~\citep{prins_toward_2015}.
To avoid the need to execute these tools manually, researchers usually write a script to combine them into what is called a workflow~\citep{perkel_workflow_2019}.

To build and maintain a complex workflow that combines many tools efficiently~\citep{leprevost_best_2014}, many workflow systems have been developed~\citep{amstutz2021existing, wratten_reproducible_2021}.
Some of these systems have large user communities, such as Galaxy~\citep{afgan_galaxy_2018}, the Common Workflow Language (CWL)~\citep{crusoe_methods_2021}, the Workflow Description Language (WDL)~\citep{voss_full-stack_2017}, Nextflow~\citep{di_tommaso_nextflow_2017}, and Snakemake~\citep{koster_snakemake_2012}.
Although each system has its unique characteristics, they have a common aim: to make computational methods portable, maintainable, reproducible, and shareable~\citep{perkel_workflow_2019}.
Most systems have a syntax for describing a workflow that is part of what is called a workflow language.
They also have an execution system that works with computational frameworks, such as a job scheduler and container virtualization~\citep{da_veiga_leprevost_biocontainers_2017}.

With the popularization of workflow systems, many research communities have worked on workflow sharing in the form of a workflow language.
Workflow registries, such as WorkflowHub~\citep{goble_implementing_2021}, Dockstore~\citep{oconnor_dockstore_2017}, and nf-core~\citep{ewels_nf-core_2020}, have been developed as public repositories for the sharing of workflows.
Workflow execution systems also utilize these registries as their tool libraries.
To improve the interoperability of workflow registries, the Global Alliance for Genomics Health (GA4GH) proposed the Tool Registry Service (TRS) specification that provides a standard protocol for sharing workflows~\citep{ga4gh_trs-api_2016, rehm_ga4gh_2021}.

Sharing workflows not only increases the transparency of research but also helps researchers by facilitating the reuse of programs, thereby making data analysis procedures more efficient.
However, workflows that are accessible on the internet are not always straightforward for others to use.
If a published workflow is not appropriately licensed, researchers cannot use it because the permission for secondary use is unclear.
A workflow may also not be executable because its format is incorrect, or dependent files cannot be found.
Even if a workflow can be executed, the correctness of its operation often cannot be verified because no tests have been provided.
Furthermore, the contact details of the person responsible for the published workflow are not always attached to it.

It is noteworthy that these issues in reusing public workflows are not often obvious to workflow developers.
To clarify the requirements for workflow sharing, Goble et al. have proposed the concept of a FAIR (findable, accessible, interoperable, and reusable) workflow~\citep{goble2020fair}.
This inheritance of the FAIR principles~\citep{wilkinson_fair_2016} focuses on the structure, forms, versioning, executability, and reuse of workflows.
However, the question remains as to who should guarantee to users that published workflows can be reused following the FAIR workflow guidelines.

WorkflowHub asks submitters to take responsibility for workflows: when a workflow is registered on WorkflowHub, the license and author identity should be clearly stated, encouraging them to publish FAIR workflows.
However, there is no obligation as to the correctness of the workflow syntax, its executability, or testing.
Not placing too many responsibilities on workflow submitters keeps obstacles to submission low, which will likely increase the diversity of public workflows on WorkflowHub; however, it will also likely increase the number of one-off submissions, which one can assume are at higher risk for the workflow problems previously described.

Unlike WorkflowHub, in nf-core, the community that operates the registry holds more accountability for published workflows.
Workflow submitters are required to join the nf-core community, develop workflows according to their guidelines, and prepare them for testing.
These requirements enable nf-core to collect workflows with remarkable reliability.
However, the community's effort tends to focus on maintaining more generic workflows that have a large number of potential users.
Consequently, nf-core states that it does not accept submissions of bespoke workflows.
This is an understandable policy, as maintaining a workflow requires domain knowledge of its content, and this is difficult to maintain in the absence of the person who developed the workflow.

In order to improve research efficiency through workflow sharing, research communities need the publication of diverse workflows in a reusable form.
However, as shown by existing workflow registries, there is a trade-off between publishing a wide variety of workflows and maintaining the reusability of the workflows that are published.
Solving this issue requires reducing the cost to developers in making and keeping their workflows reusable, which currently relies on manual effort.
This is achievable by redefining the FAIR workflow concept as a set of technical requirements and providing a system that automates their validation and testing.

We introduce Yevis, a system to share workflows with automated metadata validation and test execution.
The system expects developers and researchers who design workflows using workflow languages as users, although it does not require advanced computer skills to operate the system.
Through the development of Yevis, we specified a set of technical requirements that define a reusable workflow, according to the FAIR workflow concept.
Yevis helps researchers and communities share workflows that satisfy the requirements by supporting a build of an independent workflow registry.
To allow workflow hosting without the need of additional dedicated computing resources, Yevis works on two public web services: GitHub, a source code sharing and software development service, and Zenodo, a public research data repository.
In addition, a Yevis registry provides a web-based workflow browser and the
GA4GH TRS-compatible API ensures interoperability with other existing workflow registries.
Yevis is particularly powerful when individuals or communities want to share workflows but are without the technical expertise to build and maintain a web application.
To demonstrate that workflows can be shared that fulfill the defined requirements using Yevis, we built a registry for workflows that an existing community has managed.

\section*{Implementations}
\label{sec:implementations}

\subsection*{System design}
\label{sec:systemDesign}

\begin{figure*}[bt!]
    \centering
    \includegraphics[width=\textwidth]{figure/yevis-all-overview.pdf}
    \caption{
        The overall architecture of the yevis system.
        The registry administrator needs to set up a GitHub repository from our repository template and a Zenodo repository for file persistence.
        Workflow developers test their workflows on a local computer using our Yevis-cli, then submit a pull request to the GitHub repository.
        The GitHub repository has two GitHub Actions, testing on GitHub Actions or an external WES instance, and publishing workflow contents and metadata to the Zenodo repository and GitHub pages.
    }
    \label{fig:allOverview}
    % TODO: ref in main text
\end{figure*}

\begin{table*}
    \centering
    \vspace{\baselineskip}
    \vspace{\baselineskip}
    \caption{
        The requirements for a workflow to be considered reusable with confidence.
        We classify these requirements from the perspectives of the availability, validity, and traceability of the workflows.
        We propose that these requirements should be assured and provided to users by the workflow registries.
    }
    \label{table:wellMaintainedWorkflow}
    \begin{tabularx}{\textwidth}{llX}
        \toprule
        Perspective                   & Requirement               & Description                                                                                              \\
        \midrule
        \multirow{4}{*}{Availability} & Main workflow description & The main workflow description file is available and accessible without restriction.                      \\
                                      & Dependent materials       & The dependencies of the main workflow are available, e.g., definitions of dependent workflows and tools. \\
                                      & Testing materials         & The job configuration files for testing are available, e.g., parameter and input files.                  \\
                                      & Open-source license       & The workflow and the related materials are published under an appropriate open-source license.           \\
        \hline
        \multirow{3}{*}{Validity}     & Language type             & The language used to describe the workflow is specified, e.g., CWL, WDL, or Nextflow.                    \\
                                      & Language version          & The version of the workflow language used is specified.                                                  \\
                                      & Language syntax           & The language syntax of the workflow is valid.                                                            \\
        \hline
        \multirow{4}{*}{Traceability} & Authors and maintainers   & The contact information of the authors and the maintainers is identified.                                \\
                                      & Documentation             & The documentation of the workflow is available.                                                          \\
                                      & Workflow ID               & The unique identifier to specify the workflow is assigned, ideally by a URI.                             \\
                                      & Workflow metadata version & The version number of the workflow metadata is specified.                                                \\
        \bottomrule
    \end{tabularx}
\end{table*}

Figure~\ref{fig:allOverview} shows the overall architecture of the workflow registry built by Yevis.
The repository administrator uses our GitHub repository template and follows the guide to set up a yevis-based registry creating new repositories on GitHub and Zenodo.
After creating the metadata and passing the workflow test on a local computer, workflow developers submit the metadata as a pull request to the GitHub repository.
Once the repository receives the pull request, it automatically tests the workflow again on GitHub Actions, GitHub's continuous integration/continuous delivery (CI/CD) environment.
The system has the option to use an external WES instance for testing before accepting the submission.
The registry administrator will check the test result and approve, that is, merge the pull request.
Once the submission is approved, the repository runs another GitHub Actions automatically to upload the content to the Zenodo repository and the GitHub pages.

To implement the system, we first defined a set of requirements that the Yevis system can automatically verify or test (Table ~\ref{table:wellMaintainedWorkflow}).
By satisfying these requirements inspired by FAIR workflow, we consider a workflow is ``reusable with confidence.''
These criteria have three aspects: workflow availability, accessibility, and traceability.
The TRS defines the specification of computational tool/workflow metadata representation, including workflow's URI, used language, version, etc.
It ensures the interoperabilities among different tool/workflow registries and enables workflow engines to retrieve the information to execute a workflow maintained at a remote server, which improves the reusability of published workflows.
To help researchers share reusable workflows, we took an approach to aid them in building their own workflow registry that automatically ensures its reusability.
We define a workflow registry as a service that serves workflow information via the GA4GH TRS protocol.

The information provided by the TRS API is various workflow metadata, such as author information, documentation, language type and version, dependent materials, testing materials, etc.
Large files, such as dependent materials and testing materials, are not directly included in the TRS API response but are described as remote locations, such as HTTP protocol URLs.
Therefore, the entities that a workflow registry collects are a set of workflow metadata described in the form of the TRS API response.
In this study, therefore, we designed the system as an API server that delivers the TRS API response.

In the Yevis registry, a workflow-sharing procedure is divided into three processes: submission, review, and publication (Figure~\ref{fig:cliOverview}).
To address the requirements listed in Table~\ref{table:wellMaintainedWorkflow}, the Yevis system automatically performs processes, such as metadata validation, workflow testing, test provenance generation, persisting associated files, DOI assignment, and TRS response deployment.
To generate the TRS API response and publish it while addressing the requirements listed in Table~\ref{table:wellMaintainedWorkflow}, we implemented a command-line application called Yevis-cli.
This application contains various utilities to support the workflow registration procedure including validation and testing.
As a service and infrastructure to perform these steps, we designed Yevis to use the services of GitHub and Zenodo.
Using these web services makes it possible for communities to build a workflow registry without the need of maintaining their own computer servers.

\begin{figure*}[bt!]
    \centering
    \includegraphics[width=\textwidth]{figure/yevis-cli-overview.pdf}
    \caption{
        The flowchart of the workflow registration to a Yevis repository.
        The workflow registration procedure is divided into three processes: the submission, review, and publication process.
        Each process is performed in different locations: in the submitter's local environment, as part of the GitHub pull request, or as the GitHub Actions.
        The generated TRS API response is deployed to GitHub Pages.
        The steps indicated by yellow boxes, such as validating metadata, are performed automatically using Yevis-cli.
    }
    \label{fig:cliOverview}
\end{figure*}

\subsection*{Workflow registration with automated validation and testing}
\label{sec:buildingWorkflowRegistry}

To set up a Yevis registry, registry maintainers need to do an initial configuration of GitHub and Zenodo; this involves, for example, creating a GitHub repository, changing repository settings, and setting up security credentials.
The online documentation ``Yevis: Getting Started'' shows the step-by-step procedures to deploy a workflow registry and test it~\citep{suetake_yevis-getting-started_2022}.

We defined the Yevis metadata file, a JSON or YAML format file that contains structured workflow metadata (Figure~\ref{fig:yevisMetadataFile}).
Yevis-cli uses this file as its input and output in the submission process.
The Yevis metadata file will be published on the registry along with the TRS response to provide metadata that is not included in the TRS protocol, such as an open-source license.

\begin{figure*}[bt!]
    \centering
    \vspace{\baselineskip}
    \includegraphics[width=1.0\textwidth]{figure/yevis-metadata-file-example.png}
    \caption{
        Example of the Yevis metadata file.
        The main workflow description, dependent materials, etc. are described as remote locations; the file contains all the information that the Yevis-cli requires to automate the whole process.
        This is the actual metadata file for the workflow described in the Section ``Sharing workflows using Yevis.''
        This file is automatically updated through the processes within Yevis; for example, the file URL field is replaced by the Zenodo record URL that persists in the associated workflow files.
    }
    \label{fig:yevisMetadataFile}
\end{figure*}

\subsubsection*{Submission process}

Figure~\ref{fig:cliSubmissionProcess} shows the submission process using Yevis-cli.
During this submission process, the workflow submitter describes the workflow metadata in their local environment and submits it through a GitHub pull request (i.e., a review request to the registry maintainer).
First, Yevis-cli generates a template for the Yevis metadata file, which requires the URL of the main workflow description file as an argument.
In many workflow systems, the main workflow description file is the entry point for workflow execution.
Yevis-cli generates a template supplemented with workflow metadata automatically collected by using the GitHub REST API and inspecting the workflow's contents.
Next, the submitter needs to edit the Yevis metadata file template and add workflow tests.
Yevis-cli executes a test using a GA4GH Workflow Execution Service (WES) instance, a type of web service also described as workflow as a service~\citep{ga4gh_wes-api_2017, rehm_ga4gh_2021}; therefore, the testing materials must be written along with the specification of the WES run request.
Yevis-cli performs these tests to check if the workflow execution completes successfully.
After preparing the Yevis metadata file, Yevis-cli validates the workflow metadata syntax and runs tests using WES in the submitter's local environment.
If no WES endpoint is specified, the tests are run using Sapporo~\citep{suetake_sapporo_2022}, a production-ready implementation of WES, and Docker~\citep{merkel_docker_2014}, a container virtualization environment.
Using these portable WES environments also ensures the portability of testing in Yevis.
Finally, Yevis-cli submits the workflow as a GitHub pull request, once it confirms the required actions: the metadata validation and the test passing.
This restriction reduces the burden on the registry maintainer because many of the requirements listed in Table~\ref{table:wellMaintainedWorkflow} can be ensured during the submission process rather than the review process.

\begin{figure*}[bt!]
    \centering
    \includegraphics[width=0.85\textwidth]{figure/yevis-submission-process.pdf}
    \caption{
        The timeline of the workflow submission process using Yevis-cli.
        The submitter executes four subcommands of Yevis-cli: ``make-template,'' ``validate,'' ``test,'' and ``pull-request'' in its local environment.
        The submitter needs to edit a template of the Yevis metadata file using any text editor.
        The workflow and its metadata need to pass validation and testing before their submission, which helps to reduce the burden on the registry maintainer.
    }
    \label{fig:cliSubmissionProcess}
\end{figure*}

\subsubsection*{Review process}

Figure~\ref{fig:cliReviewProcess} shows the workflow review process using Yevis-cli.
During the review process, registry maintainers examine each workflow submitted as a Yevis metadata file on the GitHub pull request UI.
Because the submission method is restricted to Yevis-cli, the submitted workflow is guaranteed to pass validation and testing.
To ensure the reproducibility of test results on a local computer, Yevis automatically validates and tests it on GitHub Actions~\citep{suetake_yevis-test-action_2022}.
After automated validation and testing, the maintainers review the test results and log files to consider whether to approve the pull request.
Rather than using a chat tool or a mailing list, the review process through the GitHub pull request improves the transparency and traceability of workflow publication.

\begin{figure*}[bt!]
    \centering
    \vspace{\baselineskip}
    \includegraphics[width=0.85\textwidth]{figure/yevis-review-process.pdf}
    \caption{
        The timeline of the workflow review process using Yevis-cli.
        The workflow and its metadata are again validated and tested automatically on GitHub Actions.
        The test results and logs can then be reviewed by the registry maintainers with the GitHub pull request UI.
    }
    \label{fig:cliReviewProcess}
\end{figure*}

\subsubsection*{Publication process}

Figure~\ref{fig:cliPublicationProcess} shows the workflow submission process using Yevis-cli.
During the publication process, the system automatically persists all files associated with the workflow.
It generates the TRS response from the Yevis metadata file.
The approval of the pull request automatically triggers the publication process on GitHub Actions.
In the GitHub Actions script, Yevis-cli uses the Zenodo API to create a new Zenodo upload and persists all files related to the workflow~\citep{suetake_yevis-publish-action_2022}.
It obtains the DOI and persistent URLs of workflows from Zenodo, and appends them to the Yevis metadata file.
Following the Zenodo upload, the Yevis-cli in the GitHub Actions generates a TRS response JSON file and is deployed to GitHub Pages, GitHub's static web page hosting service.
Accordingly, the Yevis metadata file is merged to the default branch of the GitHub repository and deployed to GitHub Pages.
With these two files, the TRS response JSON file and the Yevis metadata file, a Yevis registry covers the information that fulfills the requirements of a reusable workflow.

\begin{figure*}[bt!]
    \centering
    \includegraphics[width=0.65\textwidth]{figure/yevis-publication-process.pdf}
    \caption{
        The timeline of the workflow publication process using Yevis-cli.
        All steps are performed automatically on GitHub Actions.
        All files related to the workflow are persisted by uploading them to Zenodo.
        The DOI is generated by Zenodo, and the Yevis metadata file is updated to append the DOI information and the persisted file URL.
        The GitHub Actions generates a TRS response from the Yevis metadata; it then deploys both of them to GitHub Pages.
    }
    \label{fig:cliPublicationProcess}
\end{figure*}

\subsection*{Workflow browsing interface}
\label{sec:resultsWorkflowBrowsingInterface}

To make it easier for registry maintainers and users to browse workflows, we implemented Yevis-web, a workflow browsing interface (Figure~\ref{fig:webOverview}).
As the interface is a browser-based application implemented in JavaScript, registry maintainers can deploy the browser on GitHub Pages.
Yevis-web accesses the TRS API served via GitHub Pages and the GitHub REST API to retrieve workflow information.
To help organize the submissions to the registry, the browser shows workflows of both statuses, those already published and those still under the review process.

\begin{figure*}[bt!]
    \centering
    \vspace{\baselineskip}
    \includegraphics[width=0.8\textwidth]{figure/yevis-web-overview.png}
    \caption{
        Screenshot of Yevis-web.
        Yevis-web is a browser-based application used via a web browser, which is deployed by workflow registry maintainers and communicates with the TRS API and GitHub REST API to retrieve workflow information.
        The browser shows both published and under-review workflows to help maintainers in organizing the registry.
        Upon selecting a workflow of interest, Yevis-web displays more detailed information, such as test results and the contents of the files related to the workflow.
    }
    \label{fig:webOverview}
\end{figure*}

\section*{Results}

\subsection*{Feature comparison with existing registries}

To clarify the advantages of a workflow registry built by Yevis, we compared the characteristics of a Yevis-based registry with WorkflowHub~\citep{goble_implementing_2021}, Dockstore~\citep{oconnor_dockstore_2017}, and nf-core~\citep{ewels_nf-core_2020}.
As comparison views, we focused on three aspects; diversity, reliability, and usability of workflows available in a registry.

In the diversity of registered workflow, as ``Acceptable workflows'' in Table~\ref{tab:featureComparison}, WorkflowHub and Dockstore have an advantage because they have no restrictions on workflows in terms of their purposes or languages.
As mentioned in the Introduction section, nf-core has the policy to collect only best-practice workflows written in Nextflow.
In contrast, a Yevis-based registry can accept any workflows written in any language as long as the registry administrator approves the submission.
The only limitation in a Yevis-based registry is the testing environment because the submission to the registry requires a suitable testing environment for the given workflow.
By default, Yevis uses Sapporo WES for its test execution, a WES implementation with multi-engine support which enables developers to extend its execution capability.

With the reliability of available workflows, we prioritize the features such as general quality control of submissions and testing preparation.
As shown in Table~\ref{tab:featureComparison}, in WorkflowHub and Dockstore, each developer is responsible for quality control and testing for the submission.
As a result, they may have workflows that are not reusable, such as those lack dependencies, documentation, or the appropriate open-source license.
The platforms do not have a strict testing policy, although it helps lower the barrier to submission.
On the other hand, nf-core does quality control and testing of its workflows by its community to provide reliable workflows.
In a Yevis-based registry, the registry itself provides automated functions to manage the quality of workflows based on the proposed requirements and test workflows in the submitter's environment and the remote CI/CD environment.

For usability, we focused on two standardized forms to identify the workflow: DOI and TRS~\ref{tab:featureComparison}.
A Yevis-based registry is only one of the four that provides DOI for each registered workflow.
Assigning DOI for workflow files prevents the problem of altering resource URLs.
For TRS compatibility, currently, nf-core is the only one not providing TRS responses.
It may be because of the design of Nextflow language, which boosts developers' productivity on a specific directory structure rather than using distributed relevant workflow files.
However, three out of four has TRS compatibility, which helps data scientist write a tool to reuse the available workflows with the unified API response.

\begin{table*}[ht!]
    \centering
    \vspace{\baselineskip}
    \vspace{\baselineskip}
    \caption{
        Feature comparison with existing registries and Yevis-based registry.
        We focused on five characteristics of registries: acceptable workflows on each registry, workflow quality control responsibility, workflow testing responsibility, DOI assignment, and TRS compatibility.
    }
    \label{tab:featureComparison}
    \begin{tabular}{lllllll}
        \toprule
        Registry       & URL             & Acceptable workflows    & Quality control by & Testing by        & DOI & TRS \\
        \midrule
        WorkflowHub    & workflowhub.eu   & No restrictions         & Each developer     & Each developer    & No  & Yes \\
        Dockstore      & dockstore.org    & No restrictions         & Each developer     & Each developer    & No  & Yes \\
        nf-core        & nf-co.re         & Generic workflows only  & nf-core community  & nf-core community & No  & No  \\
        Built by Yevis & (a GitHub repo.) & Depend on administrator & Automated by Yevis & Forced by Yevis   & Yes & Yes \\
        \bottomrule
    \end{tabular}
\end{table*}

\subsection*{Sharing workflows using Yevis}

To demonstrate that a research community can publish the workflows using Yevis while addressing the requirements listed in Table~\ref{table:wellMaintainedWorkflow}, we built a workflow registry that publishes ``DAT2-cwl'' workflows with the Yevis system~\citep{pitagora_dat2cwl_2022}\footnote{\url{https://github.com/pitagora-network/yevis-DAT2-cwl}}.
These workflows written in CWL are the appendix of the book \textit{Next Generation Sequencer DRY Analysis Manual, 2nd Edition}~\citep{bono_dat2_2019} and are maintained by the book's authors and communities.
These workflows have been maintained by a community of bioinformatics experts; however, they fulfill only a part of the requirements that we defined.
For example, the workflows have test data but would require continuous testing.
They also lack workflow metadata in a standard format.

Among the DAT2-cwl workflows, we selected a bacterial genome analysis workflow in building a new registry with Yevis~\citep{pitagora_bacteria_2022}.
This workflow combines the following command line tools: SeqKit~\citep{shen_seqkit_2016}, FastQC~\citep{simon_fastqc_2010}, fastp~\citep{chen_fastp_2018}, and Platanus-b~\citep{kajitani_efficient_2014}.
Each tool used in the workflow is packaged in a Docker container.
First, we described a Yevis metadata file (Figure~\ref{fig:yevisMetadataFile}) for this workflow using Yevis-cli and appended a test of the workflow in the form of a WES run request.
We then performed the workflow registration procedure described in the Section ``Workflow registration with automated validation and testing'' using Yevis-cli that enable the automation of many of the steps in the validation, testing, reviewing, and publishing.

Through the publication procedure of the bacteria genome analysis workflow, we evaluated how the Yevis system addressed the requirements listed in Table~\ref{table:wellMaintainedWorkflow}.
Requirements classified as ``Availability'' were addressed by being uploaded to Zenodo under an appropriate open source license~\citep{suetake_yevis-uploaded-files-bacteria_2022}.
The Yevis metadata file (Figure~\ref{fig:yevisMetadataFile})~\citep{suetake_yevis-metadata-file-bacteria_2022} and TRS API response (Figure~\ref{fig:trsApiResponse}) were updated through Yevis's publication process to use URLs persisted by Zenodo.
Requirements classified as ``Validity'' were addressed by running tests on GitHub Actions.
The contents in the Yevis metadata file and the TRS response satisfy the validity requirements, such as workflow type, workflow language version, and the URL of the test results.
Requirements classified as ``Traceability'' were addressed by describing, reviewing, and publishing them in the Yevis metadata file and TRS API response.
From the above, we confirmed that Yevis successfully published the bacteria genome analysis workflow while addressing the defined requirements.

\begin{figure*}[bt!]
    \centering
    \includegraphics[width=0.8\textwidth]{figure/api-response.pdf}
    \caption{
        TRS API response of the DAT2-cwl/bacteria-genome workflow.
        This JSON response is deployed on GitHub Pages by Yevis and is accessible via the HTTP protocol.
        The main workflow metadata in the TRS protocol is served at the path  ``/tools/\{id\}/versions/\{version\_id\}''.
        Two other possible paths for the associated files and the tests are ``/tools/\{id\}/versions/\{version\_id\}/files'' and ``/tools/\{id\}/versions/\{version\_id\}/tests''.
    }
    \label{fig:trsApiResponse}
\end{figure*}

\section*{Discussion}

Through our survey of existing workflow registries, such as Dockstore, WorkflowHub, and nf-core, it was revealed that they are maintained based on numerous contributions by various communities and the use of sufficient computer resources.
While these established workflow registries accept submissions and are available for use by researchers, there are still cases in which there is a need to create a new workflow publication platform.
For example, in the case of the Bioinformation and DDBJ center, the institute (hereafter referred to simply as DDBJ) needed to have a collection of workflows that would be allowed to run on the WES on their computing platform.
Therefore, we designed Yevis as a tool to help workflow developers create a registry to share their workflows.
DDBJ used Yevis to create and then to maintain a workflow registry dedicated to workflows for use on the DDBJ WES~\citep{suetake_ddbj-workflow-registry_2022}.

Yevis can promote the concept of a distributed workflow registry model that underlies the specifications of the GA4GH Cloud Work Stream ~\citep{rehm_ga4gh_2021}.
In the distributed workflow registry model, researchers have the option to build their own workflow registry, rather than submitting to a centralized registry.
The API standard for workflow registry specified by GA4GH enables a decentralized model, which promotes diversity in workflow development and in the research of analysis methods.
Resource sharing, particularly of analysis methods, has a bigger impact on a community studying a minor target with limited human resources.

The Yevis system strongly relies on web services, such as GitHub and Zenodo.
This is because we aimed to provide support to individuals or communities without sufficient computing resources, but this may result in a lock-in to these web services.
Although we believe these services are reliable enough to host valuable workflows, we designed the system to only use substitutable operations of those services, such as version management, file hosting, and continuous script execution.
Yet we currently provide only the implementation that depends on those two services while it is technically possible to build a Yevis registry in an on-premise server.
It is because the vendor-free system can only achieve part of our objectives to provide the effortless management of a registry.
Here, we recognize a trade-off of building an academic tool using services run by commercial vendors, which requires further discussion in the communities.
Automatic testing with GitHub Actions may also cause the issue of computational resource shortage.
To extend the capability of testing, Yevis has the option to specify the location of an external WES endpoint to run the test, which also enables the testing with a specific computational request such as GPUs or job schedulers.

Compared to existing workflow registries that have a web form for workflow registration, the Yevis system provides only a command-line interface, Yevis-cli, as a method to submit a workflow.
This is because we prefer to test workflows locally in advance of submission, while the existing registries test as part of a review process.
By using the same test suite on both the submitter's environment (local) and as part of the registry's automatic process (remote), Yevis-cli ensures better reliability of the test results.
This also helps to reduce the cost to a registry maintainer by ensuring a workflow is at least runnable on the submitter's local environment.

The Yevis system provides a well-needed solution for research communities that aim to share their workflows and wish to establish their own registry as described.
However, we recognize it still has some limitations.
One of the challenges is the description of workflow testing.
Writing tests for a workflow is difficult because the outputs may be heuristic values, may differ among tool versions, and the total amount of input and output data can be enormous.
There is a need for a testing framework that packages test cases and results and generates assertions that allow for slight differences in the output.
Debugging a workflow is also difficult because a typical workflow uses many tools that use various programming runtimes.
Therefore, a framework is required to capture metrics of the test execution environment with these various runtimes.

Another challenge for the proposed distributed registry model is the findability of workflows.
In the model where each developer is responsible for their content, the use of appropriate terms for describing workflow metadata can be an issue.
A possible solution to improve the findability of workflows in distributed registries is to collect metadata in a centralized registry to curate them and create the search index.
However, this will require a further challenge to distinguish the collected workflows using only metadata.

Many researchers agree that resource sharing is a key factor in the era of data science.
As workflow systems and their communities grow, researchers have worked to share their data analysis procedures along with their data.
Despite the fact that workflow systems are developed for automation, it sounds strange that maintaining workflow registries still relies on manual efforts.
Through the development of Yevis, we found there are many possibilities for further automation in the process of resource sharing.
Through the defined requirements for reusable workflows and a system that ensures them automatically, we believe that our work can contribute to moving open science forward.

\section*{Availability of source code and requirements}

\begin{itemize}
    \setlength{\itemsep}{0.5mm}
    \item Project name: Yevis-cli
    \item Project home page: \url{https://github.com/ddbj/yevis-cli}
    \item DOI: 10.5281/zenodo.6541109
    \item biotoolsID: yevis-cli
    \item Operating system(s): Platform independent
    \item Programming language: Rust
    \item Other requirements: Docker recommended
    \item License: Apache License, Version 2.0
\end{itemize}

\vspace{2mm}
\begin{itemize}
    \setlength{\itemsep}{0.5mm}
    \item Project name: Yevis-web
    \item Project home page: \url{https://github.com/ddbj/yevis-web}
    \item DOI: 10.5281/zenodo.6541031
    \item biotoolsID: yevis-web
    \item Operating system(s): Platform independent
    \item Programming language: TypeScript
    \item License: Apache License, Version 2.0
\end{itemize}

\section*{Availability of supporting data and materials}

Data and materials related to the DAT2-cwl workflows described in the Section ``Sharing workflows using Yevis'' are available on GitHub and Zenodo as follows:

\begin{itemize}
    \setlength{\itemsep}{0.5mm}
    \item GitHub repository for DAT2-cwl workflows~\citep{pitagora_dat2cwl_2022}
    \item Workflow registry yevis-DAT2-cwl~\citep{suetake_yevis-dat2-cwl_2022}
    \item Workflow browser for yevis-DAT2-cwl~\citep{suetake_yevis-dat2-cwl-browser_2022}
\end{itemize}

\section*{Declarations}

\subsection*{List of abbreviations}

API: Application Programming Interface;
CI/CD: Continuous Integration/Continuous Delivery;
CWL: Common Workflow Language;
DDBJ: Bioinformation and DDBJ Center;
DNA: Deoxyribonucleic Acid;
DOI: Digital Object Identifier;
FAIR: Findable, Accessible, Interoperable, and Reusable;
GA4GH: Global Alliance for Genomics and Health;
HTTP: Hypertext Transfer Protocol;
ID: Identifier;
REST: Representational State Transfer;
TRS: Tool Registry Service;
UI: User Interface;
URI: Uniform Resource Identifier;
URL: Uniform Resource Locator;
WDL: Workflow Description Language;
WES: Workflow Execution Service;

\subsection*{Ethical Approval}

Not applicable for this study.

\subsection*{Consent for publication}

Not applicable for this study.

\subsection*{Competing Interests}

The authors declare that they have no competing interests.

\subsection*{Funding}

This study was supported by JSPS KAKENHI Grant Number 20J22439, the Life Science Database Integration Project, and the National Bioscience Database Center (NBDC) of the Japan Science and Technology Agency (JST).
This study was also supported by the CREST program of the Japan Science and Technology Agency (Grant Number JPMJCR17A1).

\subsection*{Author's Contributions}

H.S. and T.O. conceived and developed the methodology and software and conducted the investigation. H.S., T.F., and T.O. wrote the manuscript. T.F., T.I., and T.O. supervised the project. All authors read and approved the final version of the manuscript.

\section*{Acknowledgements}

We acknowledge and thank the following scientific communities and their collaborative events where several of the authors engaged in irreplaceable discussions and development throughout the project: the Pitagora Meetup, Workflow Meetup Japan, NBDC/DBCLS BioHackathon Series, and Elixir's BioHackathon Europe Series.
We also would like to thank Ascade Inc. for their support with the software development.

\section*{References}

\bibliography{main}

\end{document}
