## Supplementary figures and images for "Workflow sharing with automated metadata validation and test execution to improve the reusability of published workflows"

### yevis-all-overview.pdf

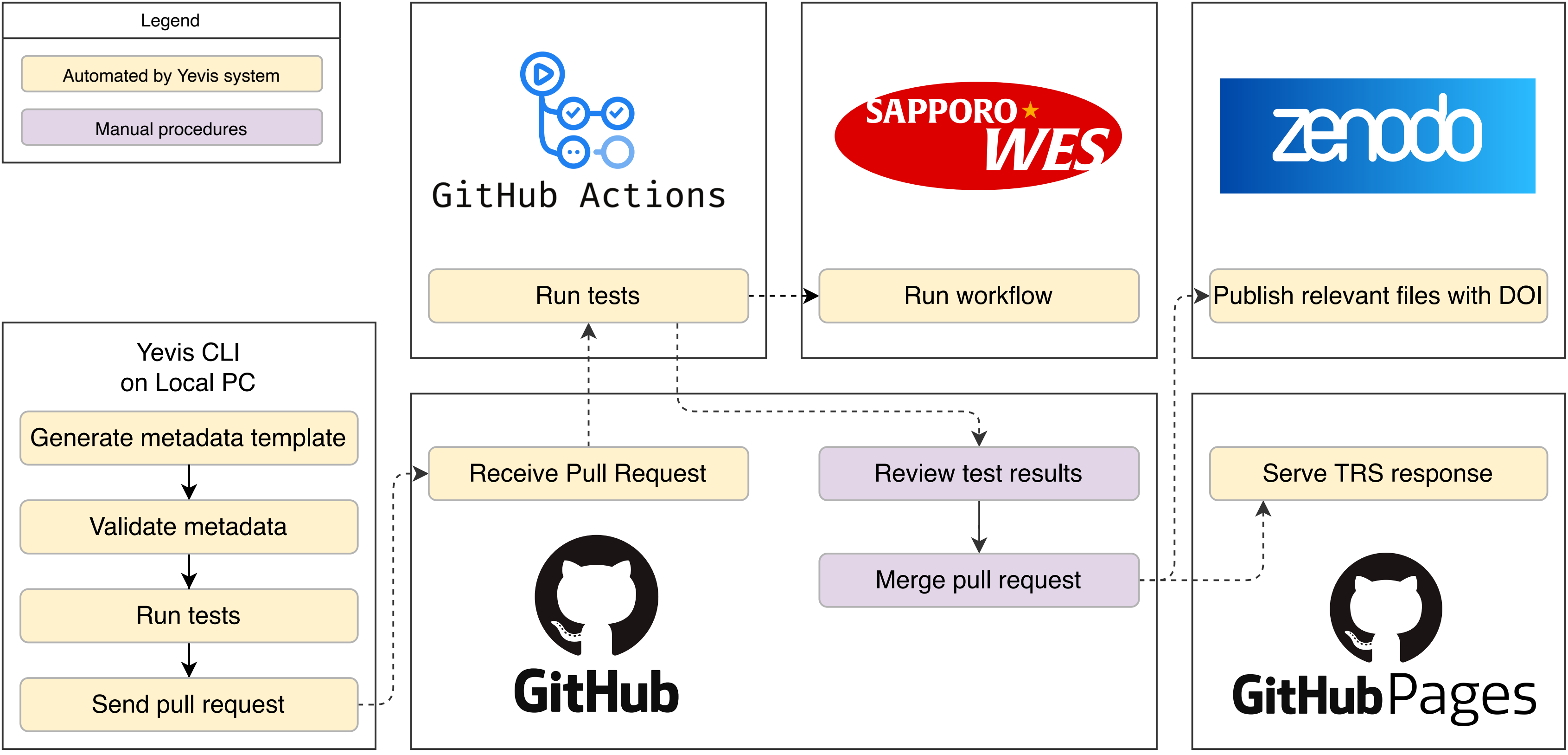

### yevis-cli-overview.pdf

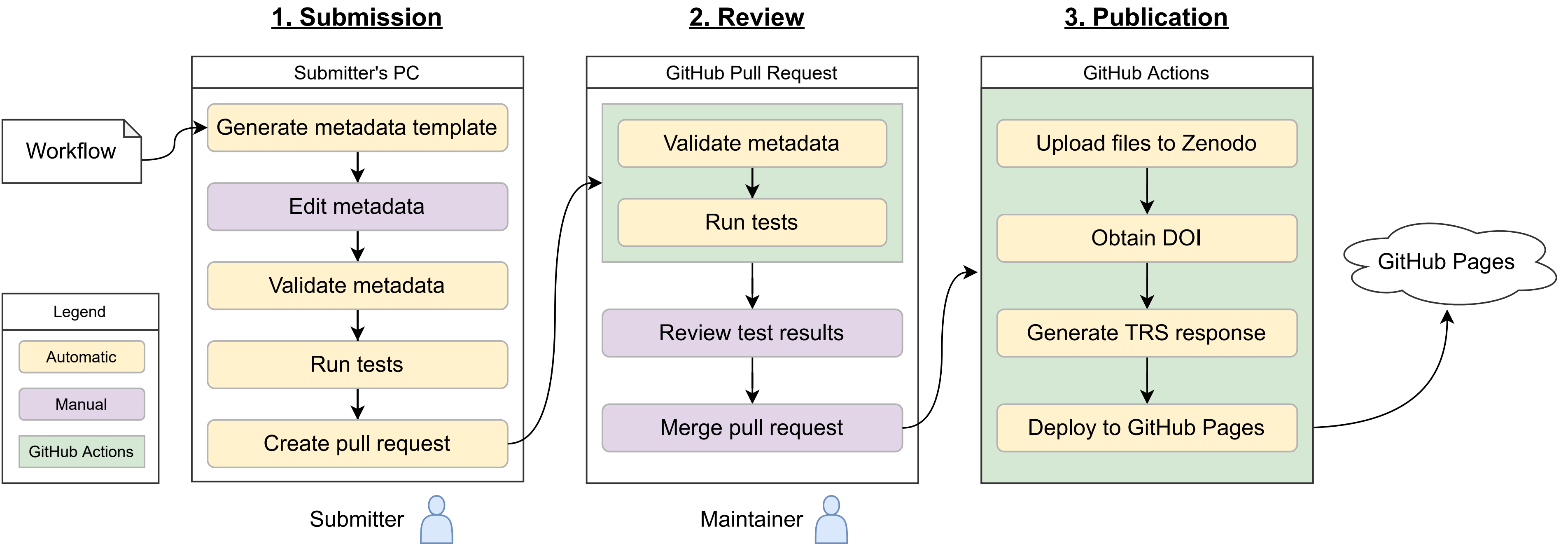

### yevis-metadata-file-example.png

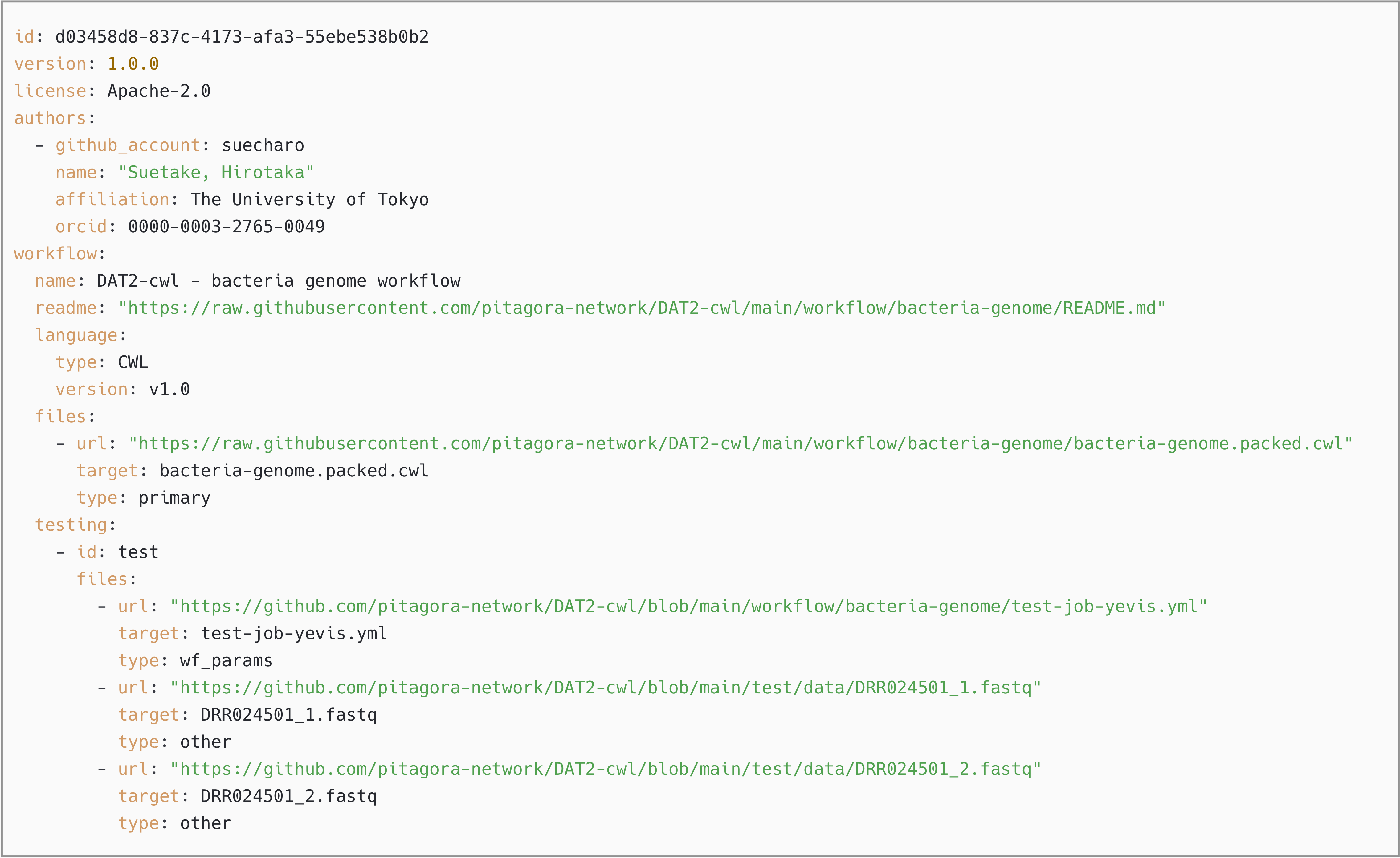

### yevis-publication-process.pdf

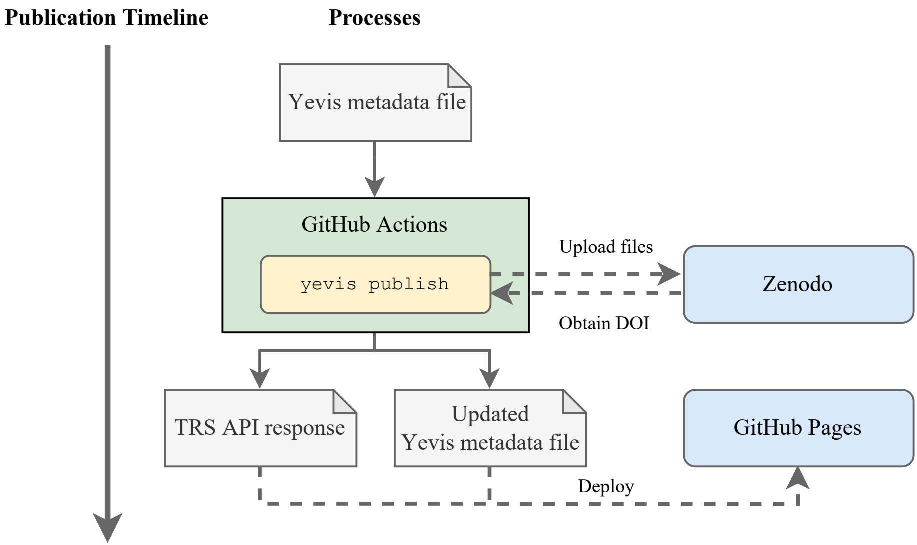

### yevis-review-process.pdf

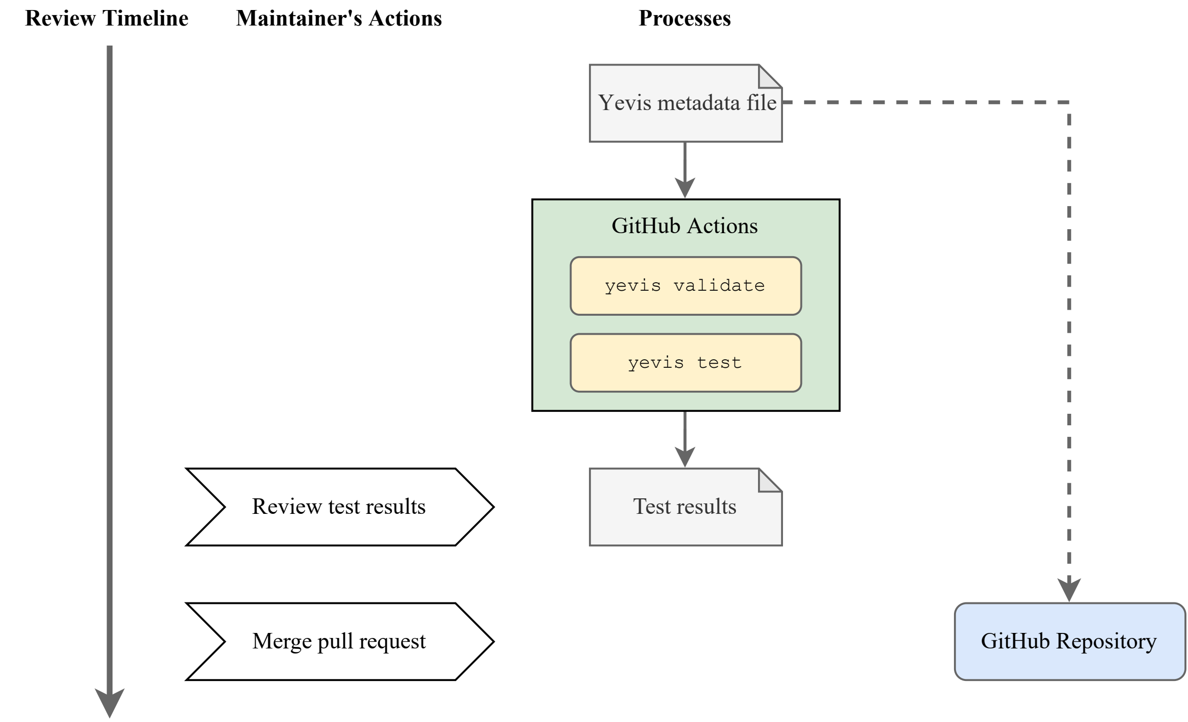

### yevis-submission-process.pdf

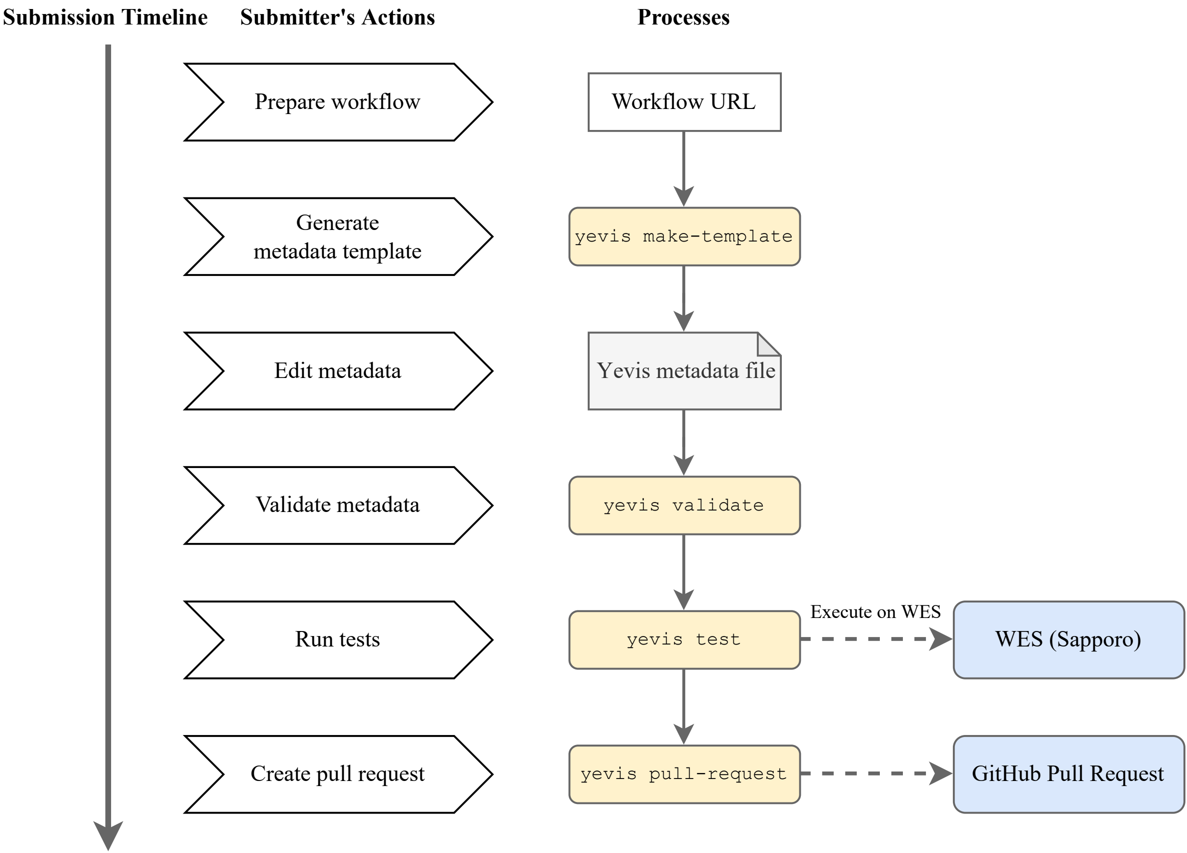

### yevis-web-overview.png

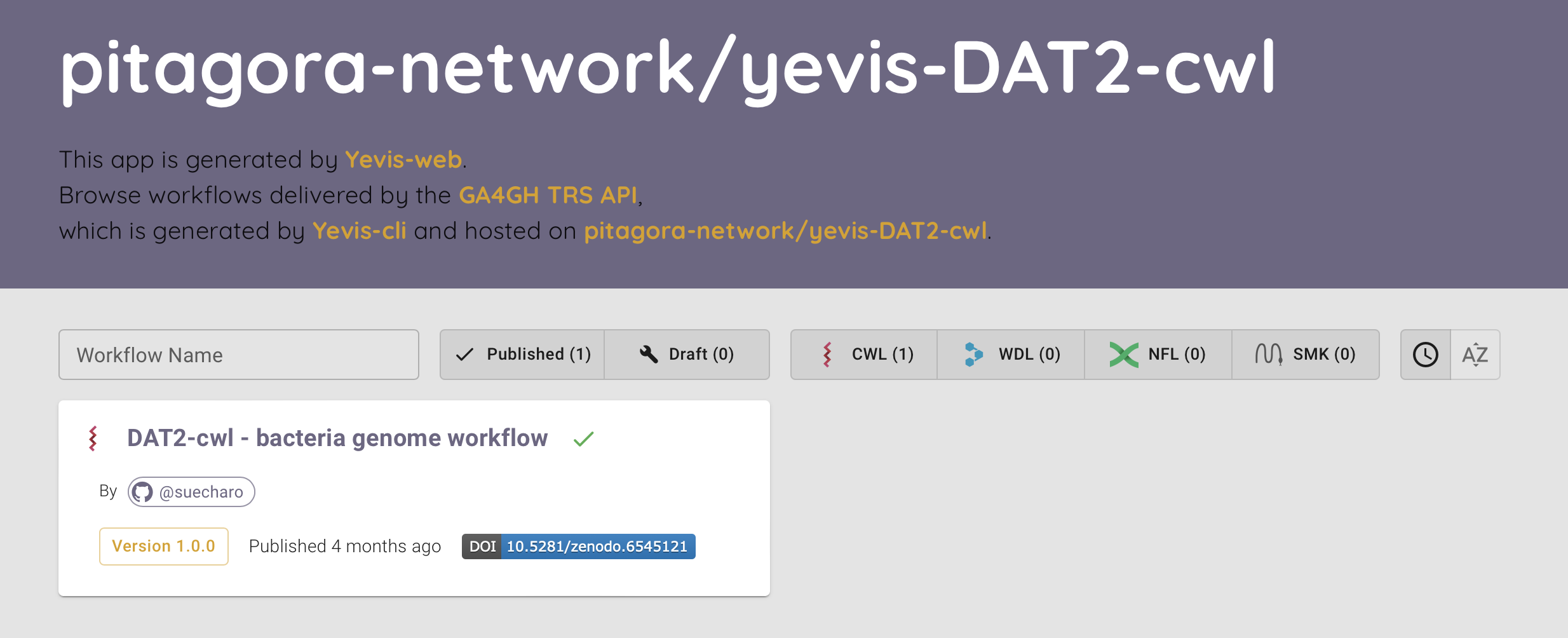
